# Gene editing and mutagenesis reveal inter-cultivar differences and additivity in the contribution of *TaGW2* homoeologues to grain size and weight in wheat

**DOI:** 10.1101/327874

**Authors:** wei wang, James Simmonds, Qianli Pan, Dwight Davidson, Fei He, Abdulhamit Battal, Alina Akhunova, Harold N. Trick, Cristobal Uauy, Eduard Akhunov

## Abstract

The *TaGW2* gene homoeologues have been reported to be negative regulators of grain size (GS) and thousand grain weight (TGW) in wheat. However, the contribution of each homoeologue to trait variation among different wheat cultivars is not well documented. We used the CRISPR-Cas9 system and TILLING to mutagenize each homoeologous gene copy in cultivars Bobwhite and Paragon, respectively. Plants carrying single-copy nonsense mutations in different genomes showed different levels of GS/TGW increase, with TGW increasing by an average of 5.5% (edited lines) and 5.3% (TILLING mutants). In any combination, the double homoeologue mutants showed higher phenotypic effects than the respective single-genome mutants. The double mutants had on average 12.1% (edited) and 10.5% (TILLING) higher TGW with respect to wild-type lines. The highest increase in GS and TGW was shown for triple mutants of both cultivars, with increases of 16.3% (edited) and 20.7% (TILLING) in TGW. The additive effects of the *TaGW2* homoeologues were also demonstrated by the negative correlation between the functional gene copy number and GS/TGW in Bobwhite mutants and an F_2_ population. The highest single-genome increases in GS and TGW in Paragon and Bobwhite were obtained by mutations in the B and D genomes, respectively. These inter-cultivar differences in the phenotypic effects between the *TaGW2* gene homoeologues coincide with inter-cultivar differences in the homoeologue expression levels. These results indicate that GS/TGW variation in wheat can be modulated by the dosage of homoeologous genes with inter-cultivar differences in the magnitude of the individual homoeologue effects.

## Introduction

Wheat is one of the most widely grown crops in the world and contributes 20% of the calories in human diets (FAO: http://www.fao.org/faostat/). To ensure food security and to achieve sustainable increases in wheat production, novel breeding strategies underpinned by the molecular understanding of traits contributing to yield need to be developed and deployed. The identification of genes controlling yield component traits in many crops (Li and Yang 2017) and the development of novel wheat genomic resources and reverse genetics tools (Krasileva et al. 2017; Wang et al. 2018) provide unique opportunities to improve yield potential in wheat.

The allohexaploid genome of common wheat (*Triticum aestivum*, 2n = 6x, genomes *AABBDD*) has many genes present in three copies on homoeologous chromosomes. This genetic redundancy impedes the functional studies of genes controlling important agronomic traits, because the phenotypic effects of loss-of-function mutations in a single homoeologue are frequently masked by other gene copies (Borrill et al. 2015). The development of reverse genetics methods based on either TLLING (Krasileva et al. 2017) or CRISPR-Cas9 gene editing (Wang et al. 2016; Wang et al. 2018) provide powerful tools to understand the role of wheat gene homoeologues in controlling complex phenotypic traits and to expand genetic diversity accessible for wheat improvement offsetting the negative effects of domestication and improvement bottlenecks on genetic variation (Avni et al. 2017; Cavanagh et al. 2013; Wang et al. 2014a).

The *TaGW2* gene is an orthologue of the rice *GW2* gene which negatively regulates grain width and TGW (thousand grain weight) by modulating cell division on the husk and grain filling (Song et al. 2007). The A genome homoeologue of the *TaGW2* gene (*TaGW2-A1*) was previously validated as a negative regulator of grain width, grain length and TGW in wheat by forward genetic methods (Jaiswal et al. 2015; Su et al. 2011; Yang et al. 2012). Using the complementary reverse genetics approach, a tetraploid wheat mutant of *TaGW2-A1* was identified from a TILLING population (Uauy et al. 2009) and the phenotypic effects of *TaGW2-A1* was studied by backcrossing the mutant line with tetraploid wheat Kronos and hexaploid wheat Paragon (Simmonds et al. 2016). The results showed that the knock-out (KO) of *TaGW2-A1* increased grain width (2.8%), grain length (2.1%) and TGW (6.6%) in both tetraploid and hexaploid wheat, further validating *TaGW2-A1* as a negative regulator of GS and TGW.

The CRISPR-Cas9 based genome editing technology has recently been applied widely to study gene function in many species (Cong et al. 2013; Jinek et al. 2012; Mali et al. 2013; Minkenberg et al. 2017; Mohan 2016; Wang et al. 2018). The CRISPR-Cas9 system has been successfully used in wheat to produce mutant plants for several genes (Liang et al. 2017; Wang et al. 2018; Wang et al. 2014b; Zhang et al. 2016). We previously showed that the KO of all three homoeologues of *TaGW2* gene by CRISPR-Cas9 increased GS and TGW much higher than the reported increase of *TaGW2-A1* mutants (TGW; 28% compared to 7%), which suggests that the B and D homoeologues (*TaGW2-B1* and *TaGW2-D1*) have potential phenotypic effects (Wang et al. 2018). Recently, the B and D homoeologues were also shown to contribute to GS (grain size) and TGW in an additive manner; a double mutant line with mutations in the B and D homoeologues produced stronger phenotypic effects than single mutant lines for either the B or D homoeologues (Zhang et al. 2018). However, these studies did not fully uncover the genetic relationship among all three *TaGW2* homoeologues.

In this study, we used the sequenced wheat TILLING populations (Krasileva et al. 2017) and CRISPR-Cas9-based genome editing to investigate the contribution of the *TaGW2-A1*, -*B1* and - *D1* gene homoeologues to phenotypic variation in GS and TGW in two hexaploid wheat cultivars. Plants carrying mutations in *TaGW2-A1*, -*B1* and -*D1* in all possible combinations were obtained in transgenic cultivar Bobwhite by expressing the CRISPR-Cas9 gene editing plasmids and in cultivar Paragon by crossing mutants identified via conventional (Uauy et al. 2009) and the sequenced (Krasileva et al. 2017) TILLING population. The increases of GS and TGW obtained for the triple- and the three double-mutant combinations were additive with respect to single mutants. This negative correlation between *TaGW2* functional allele numbers (6 in total) and GS/TGW was also validated in an F_2_ population. We found inter-cultivar differences in the contribution of different genomes to the GS and TGW phenotypes; mutations in *TaGW2-D1* and *TaGW2-B1* explained most of the phenotypic variation in Bobwhite and Paragon, respectively. These differences appear to be associated with the level of the homoeologue expression relative to other gene copies, consistent with variation in the expression of *TaGW2* homoeologues among different wheat cultivars. Our results suggest that in addition to dosage-dependent regulation of the GS and TGW traits, different homoeologues copies of *TaGW2* may play a role in defining the magnitude of phenotypic effects for these traits in different wheat cultivars.

## Materials and Methods

### CRISPR-Cas9-induced *TaGW2* gene mutant plants

The T_0_ generation plants carrying CRISPR-Cas9-induced mutations were regenerated and genotyped as described previously (Wang et al. 2018). Briefly, the CRISPR-Cas9 targeted region on the first exon of *TaGW2-A1, -B1* and -*D1* was amplified by PCR using gene-specific primers with 5’-tails including part of the Illumina Truseq adapters. The second round of PCR was used to add the Illumina Truseq adapter barcodes. To expand the multiplexing capacity of Illumina TruSeq barcodes, additional five barcoding bases were added between the target-specific primers and Illumina adaptors. As shown in Table S1, 96 of samples carrying the same TruSeq barcode could be indexed using these internal barcodes.

The T_1_, T_2_ or T_3_ generation progeny from independent T_0_ lines were used in phenotyping experiments. Cultivar Bobwhite and plants carrying the wild-type alleles of *TaGW2* genes segregating from the same T_0_ lines were used as controls. A T_0_ *TaGW2* triple genome mutant was crossed with cultivar Thatcher. The F_1_ plants were self-pollinated and seeds grown to produce F_2_ populations. Because the transgenerational activity of CRISPR-Cas9 constructs could induce new mutations in the non-edited target sites (Wang et al. 2018), all plants in different experiments were genotyped as described above (the mutations are shown in Table S2 - S4). Any plant carrying deletions or insertions not causing coding frame shift were excluded from further analyses.

### Selection EMS-induced loss-of-function mutants of the *TaGW2* gene

The homozygous G to A mutation at the AG splice acceptor site of exon 5 (G2373A) in *TaGW2-A1* was discovered through screening of the Kronos TILLING population as reported in Simmonds et. al (2016).

Primers specific to the B genome (JB2 & JB7) were designed to amplify a region of 1091-bp across exons 2-6 of *TaGW2-B1* (Table S5). The first three pooled DNA plates of the Kronos TILLING population were screened using the same procedures as for the A genome. In total 19 putative mutations were discovered, one resulting in a heterozygous C to T transition at position 514 of the *TaGW2-B1* coding sequence (CDS) in exon 5 (position 2504 in gDNA), leading to a premature stop codon in TILLING line Kronos0341. To confirm the mutation, a KASP (Kompetitive Allele Specific PCR) marker was developed (*TaGW2_B1*_WT/M/C) to genotype the C/T polymorphism. The common primer utilizes a SNP unique to the B genome for specificity.

The D genome mutation was discovered through the in-silico wheat TILLING database (www.wheat-tilling.com) (Krasileva et al. 2017) using a BLASTN-based comparison of the *TaGW2-D1* sequence. TILLING line Cadenza1441 was identified as containing a G to A transition at position 698 of CDS in exon 7 (position 7139 in gDNA) causing a premature termination codon. The pre-designed D genome specific KASPar marker available on the website (Ramirez-Gonzalez et al. 2015) was utilized to confirm the mutation. The KASPar markers designed to track the mutations are listed in Table S5. The locations of the mutations for the three homoeologues and the resulting amino acid changes are shown in Figure S1.

### Development of wheat lines with double and triple EMS KOs of the *TaGW2* gene

TILLING mutant Kronos0341, which carries the heterozygous C2504T mutation in *TaGW2-B1*, was crossed to the cultivar Paragon. Plants of TILLING mutant Cadenza1441 carrying the homozygous mutation G7139A in *TaGW2-D1* were crossed with Paragon NILs homozygous for the G2373A *TaGW2-A1* SNP. The resulting F_1_ plants were inter-crossed to produce BC_1_F_1_ seeds (Figure S2). Marker assisted selection was carried out on the BC_1_F_1_ plants using KASPar markers for the selection of plants heterozygous for all three mutations (*AaBbDd*). These plants were self-pollinated and the resultant plants (BC_1_F_2_) screened for the selection of lines homozygous for all three mutations. To enable a replicated experiment plants were once more self-pollinated to BC_1_F_3_.

Seeds of the Paragon BC_1_ NIL carrying the three TILLING derived mutant alleles developed in this study are available to the wheat community via the JIC Germplasm Resources Unit (https://seedstor.ac.uk/) accession number W10347.

### Plant growth conditions

The CRISPR-Cas9 mutant plants were grown in Kansas State University greenhouses under 12/12 hour light/dark conditions in the 1^st^ month, and then grown until harvested under 16 hour light/8 h dark; room temperature was set as 24 °C in the day and 21 °C in the night. Plants were grown in 1 liter square pots filled with 3/4 lab made soil (volume ratio soil:peatmoss:perlites:CaSO_4_ is 20:20:10:1) on the bottom and 1/4 Sungro soil (Sun Gro Horticulture, Agawam, MA, USA) on the top. The F_2_ populations of Bobwhite’s *TaGW2* gene mutant and Thatcher were grown in the 1/4 liter pots under the same greenhouse conditions as Bobwhite mutant plants. For each experiment, all plants were set in one greenhouse randomly.

The TILLING derived BC_1_F_3_ plants were sown at the John Innes Centre in two 96 well trays comprising of peat and sand compost. The seedlings were propagated for three weeks in a controlled environment room (CER) with set points of 16 hours day/8 hours night and temperatures of 20/16 °C. Once genotyped, selected plants were potted into 1 liter pots with Petersfield Cereal Mix (Petersfield, Leicester, UK) and transferred to a Conviron BDW80 CER (Conviron, Winnipeg, Canada) set at 16/8 day/night lighting (300 μmol m^−2^ s^−1^), temperatures of 20/15°C respectively and 50% relative humidity, until maturity. Plants were arranged in a complete randomized design.

### Expression analysis of *TaGW2* gene

To estimate the relative gene expression levels of *TaGW2-A1*, -*B1* and -*D1* in Bobwhite, the tissues from pollinating stage Bobwhite plants were sampled and frozen immediately in liquid nitrogen. In total, tissues from five plants were collected. Seven different tissues were sampled from the main spike including one week-old endosperm, spike-removed endosperm, flag leaf, flag leaf sheath, leaf and leaf sheath (two leaves lower than flag leaf), and stem. In addition, one non-pollinated spike and roots were sampled from the same plants. RNA was extracted using Trizole (Thermo Fisher Scientific, Catalog #: 15596018) following the manufacture’s protocol. The first-strand cDNA was synthesized using kit “SuperScript™ III First-Strand Synthesis SuperMix for qRT-PCR” (Thermo Fisher Scientific, Catalog #: 11752-250) following the manufacture’s protocol. The *TaGW2-A1*, -*B1* and -*D1* homoeologue-specific primers (Table S6) were designed and validated by PCR using the DNA of Chinese Spring nulli-tetrasomic lines (Figure S3). Real-time PCR was performed with iQ™ SYBR Green Supermix (BIO-RAD, Catalog #: 170-8882) following the manufacture’s protocol, using the *TaActin* gene as reference.

The relative expression levels of the *TaGW2-A1*, -*B1* and -*D1* genes in different cultivars were assessed in one week-old seedlings. The relative expression level of *TaGW2-A1*, -*B1* and -*D1* in the Bobwhite lines carrying different mutated alleles of *TaGW2* were analyzed in the 5^th^ leaf of plants at the 5-leaf development stage.

The relative expression levels of *TaGW2-A1*, -*B1* and -*D1* in hexaploid cultivar Azhurnaya was downloaded from the wheat eFP browser (http://bar.utoronto.ca/efp_wheat/cgi-bin/efpWeb.cgi; Ramirez-Gonzalez et al. *under review*) based on methods outlined in Borrill et al (Borrill et al. 2016). The expression levels of the *TaGW2-A1*, -*B1* and -*D1* genes in cultivar Chinese Spring were obtained from WheatExp (https://wheat.pw.usda.gov/WheatExp/) (Pearce et al. 2015) and were based on RNA-Seq data generated for tissues at different developmental stages defined by Zadoks scale (Choulet et al. 2014; Zadoks et al. 1974).

### Grain morphometric and TGW trait data

A MARVIN seed analyzer (GTA Sensorik GmbH, Germany) was used to collect data on grain morphometric measurements (grain width, length, area), and thousand grain weight. All seeds on a plant from the CRISPR-Cas9 mutant populations were analyzed, and the mean values per plant were used for statistical analyses. In Paragon mutants, seed data was collected from three spikes per plant separately, and the mean of three spikes were calculated and used for further analyses.

### Statistical analysis of data

The distribution of raw GS/TGW data was visualized using the box and whisker plots. One way ANOVA was applied when multiple groups of data were compared followed by post-hoc Tukey’s test. The Student’s t-test was applied to assess the significance of difference between the two groups of data.

## Results

### Single genome KO mutants of *TaGW2* homoeologues increase both grain size and weight

The phenotypic effects of a KO mutation in only one of the three *TaGW2* gene homoeologues were assessed by comparing grain morphometric parameters and TGW of mutant and wild type wheat lines. In Bobwhite, all KO mutations in the *TaGW2-A1*, *TaGW2-B1* or *TaGW2-D1* gene copies significantly increased the grain width, grain area and TGW in at least one experiment (Student’s t-test *P* < 0.05; Table 1 and 2, Figure 1 and S4). The KO mutation of *TaGW2-D1* gene had the highest effects, increased the grain width, grain area and TGW by 2.3 ~ 2.9%, 4.6 ~ 5.9% and 6.7 ~ 8.0%, respectively, while the KO mutation of *TaGW2-A1* gene had the lowest effects, increased the grain width, grain area and TGW by 1.7 ~ 2.2%, 0.2 ~ 3.7% and 3.9 ~ 4.9%, respectively. The increase in TGW for single mutants had a weighted average of 5.46% across all single mutants. The grain length was significantly increased only in the KO mutant of the *TaGW2-D1* gene (3.0 ~ 3.8%), but not the KOs of the *TaGW2-A1* and *TaGW2-B1* genes. The comparison of the seed morphometric traits and TGW between the regenerated wheat lines and Bobwhite cultivar revealed no differences indicating that plant transformation and regeneration did not affect the traits of interest in gene edited plants (Table 2).

**Figure 1.**
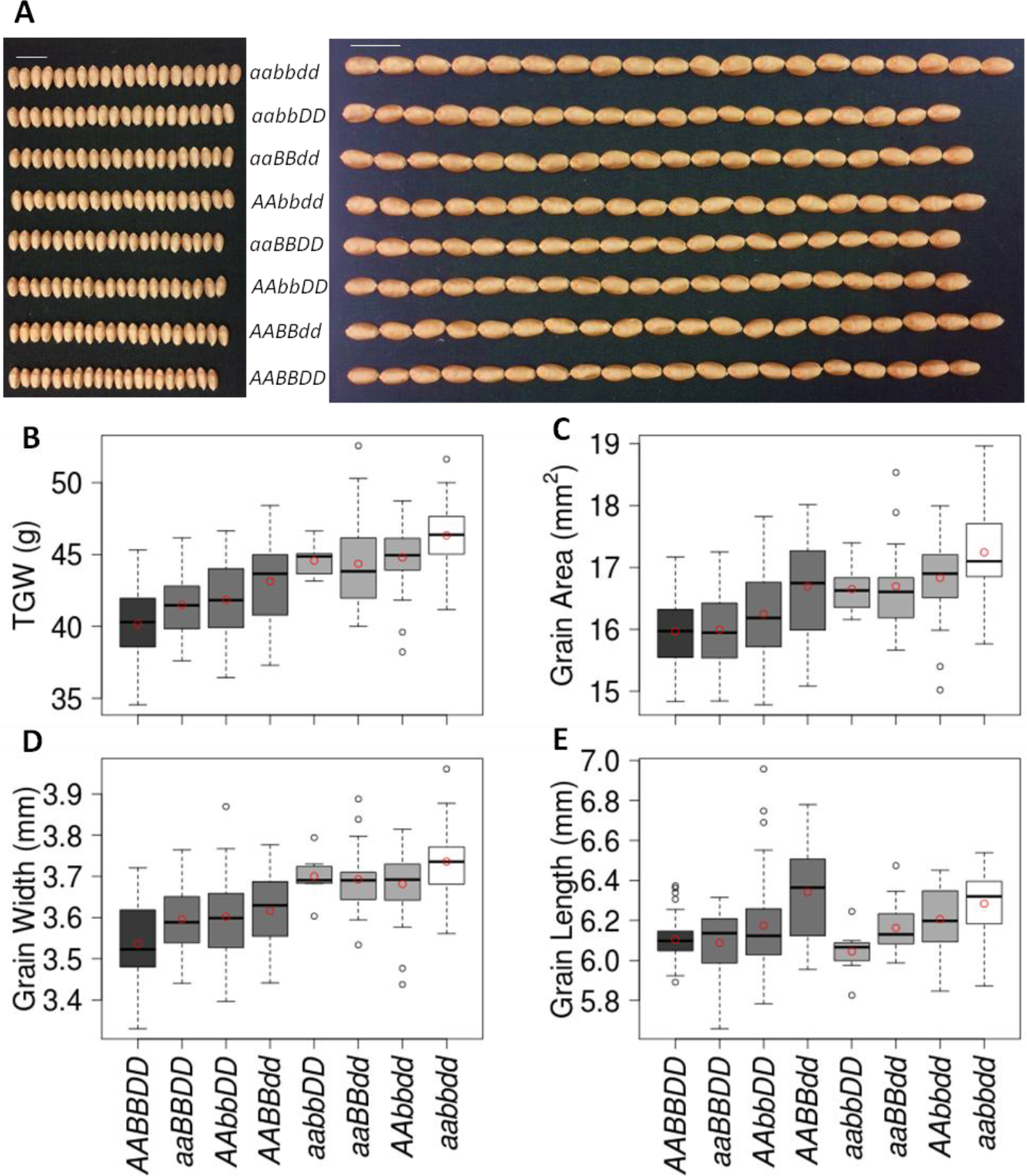
The effects of single-, double-, and triple- KO mutations in the *TaGW2* gene homoeologues on the grain morphometric and TGW traits in Bobwhite. **A)** The image of twenty seeds from wild-type, single-, double- and triple- mutant plants (scale bar 1 cm). **B-E**) Box and whisker plots show the distribution of TGW (B), grain area (C), grain width (D), and grain length (E) for wild-type and mutant wheat lines. The datasets from Bobwhite and the T_0_ progeny plants carrying wild-type *TaGW2* alleles were combined because they did not show statistical differences (Table 2). The mean value for each genotype is shown as a red circle. The genotypes of the *TaGW2* homoeologues are shown in all panels with lower and uppercase letters corresponding to the mutant and wild-type alleles, respectively, for the A, B, and D genome homoeologues.

**Table 1.**
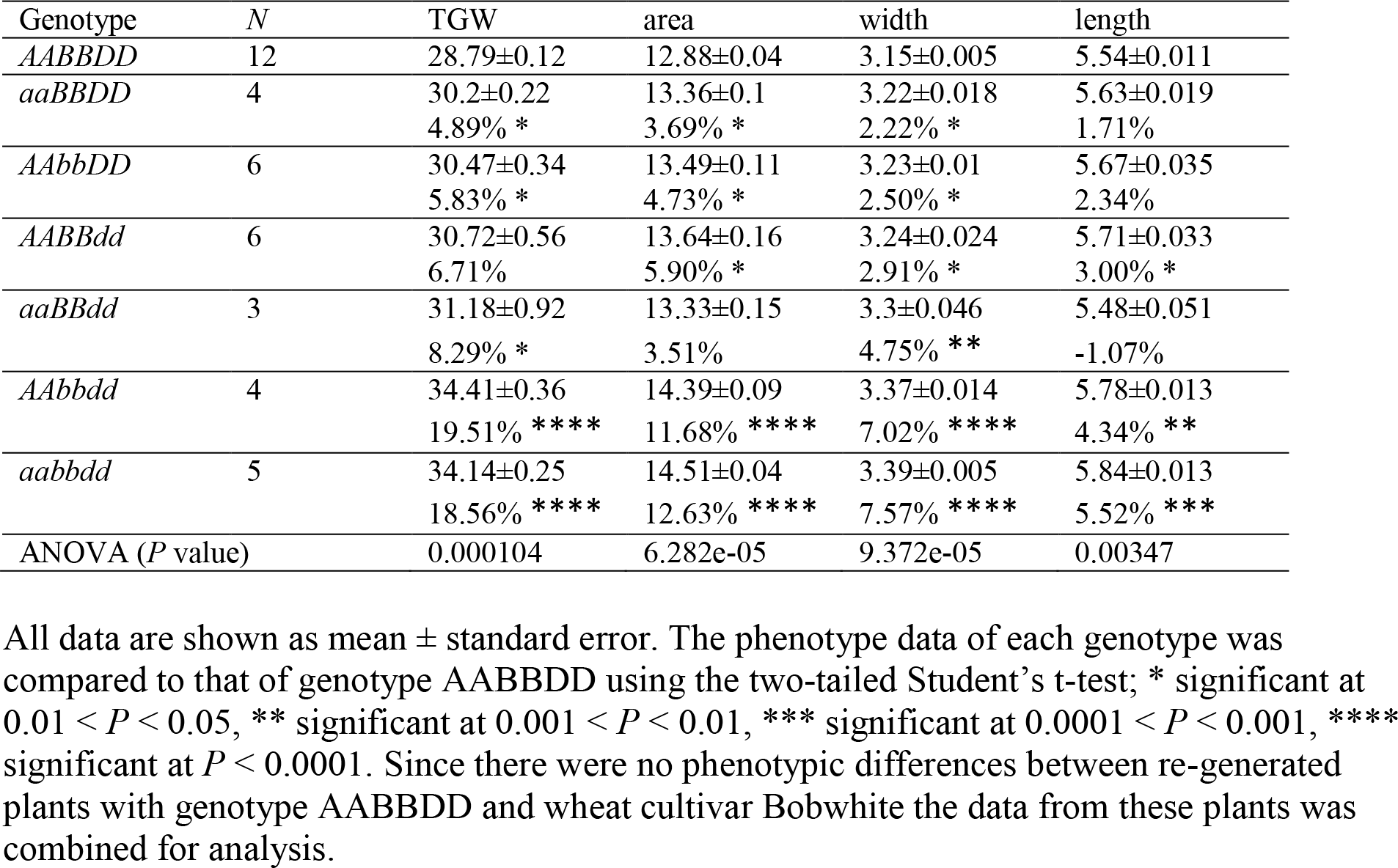
The thousand grain weight and grain morphometric parameters for CRISPR-Cas9-induced *TaGW2* gene mutants (2017 Spring experiment).

| Genotype | *N* | TGW | area | width | length |
| --- | --- | --- | --- | --- | --- |
| *AABBDD* | 12 | 28.79±0.12 | 12.88±0.04 | 3.15±0.005 | 5.54±0.011 |
| *aaBBDD* | 4 | 30.2±0.22<br>4.89% * | 13.36±0.1<br>3.69% * | 3.22±0.018<br>2.22% * | 5.63±0.019<br>1.71% |
| *AAbbDD* | 6 | 30.47±0.34<br>5.83% * | 13.49±0.11<br>4.73% * | 3.23±0.01<br>2.50% * | 5.67±0.035<br>2.34% |
| *AABBdd* | 6 | 30.72±0.56<br>6.71% | 13.64±0.16<br>5.90% * | 3.24±0.024<br>2.91% * | 5.71±0.033<br>3.00% * |
| *aaBBdd* | 3 | 31.18±0.92<br>8.29% * | 13.33±0.15<br>3.51% | 3.3±0.046<br>4.75% ** | 5.48±0.051<br>-1.07% |
| *AAbbdd* | 4 | 34.41±0.36<br>19.51% **** | 14.39±0.09<br>11.68% **** | 3.37±0.014<br>7.02% **** | 5.78±0.013<br>4.34% ** |
| *aabbdd* | 5 | 34.14±0.25<br>18.56% **** | 14.51±0.04<br>12.63% **** | 3.39±0.005<br>7.57% **** | 5.84±0.013<br>5.52% *** |
| ANOVA (*P* value) |  | 0.000104 | 6.282e-05 | 9.372e-05 | 0.00347 |

All data are shown as mean ± standard error. The phenotype data of each genotype was compared to that of genotype *AABBDD* using the two-tailed Student's t-test; \* significant at $0.01 < P < 0.05$ , \*\* significant at $0.001 < P < 0.01$ , \*\*\* significant at $0.0001 < P < 0.001$ , \*\*\*\* significant at $P < 0.0001$ . Since there were no phenotypic differences between re-generated plants with genotype *AABBDD* and wheat cultivar Bobwhite the data from these plants was combined for analysis.

**Table 2.** The thousand grain weight and grain morphometric parameters for CRISPR-Cas9-induced *TaGW2* gene mutants (2017 Fall experiment).

| Genotype | N | TGW(g) | Area(mm<sup>2</sup>) | Width(mm) | Length(mm) |
| --- | --- | --- | --- | --- | --- |
| Bobwhite | 11 | 40.85±0.53 | 15.97±0.14 | 3.53±0.018 | 6.1±0.038 |
| *AABBDD* | 37 | 39.95±0.16 | 15.96±0.08 | 3.54±0.056 | 6.11±0.029 |
| *aaBBDD* | 26 | 41.49±0.46<br>3.85% * | 15.99±0.13<br>0.19% | 3.6±0.017<br>1.69% * | 6.09±0.032<br>-0.33% |
| *AAbbDD* | 53 | 41.85±0.37<br>4.76% ** | 16.25±0.11<br>1.82% | 3.6±0.013<br>1.69% ** | 6.17±0.032<br>0.98% |
| *AABBdd* | 28 | 43.15±0.56<br>8.01% **** | 16.69±0.16<br>4.57% *** | 3.62±0.017<br>2.26% ** | 6.34±0.045<br>3.76% **** |
| *aabbDD* | 7 | 44.58±0.45<br>11.59% *** | 16.65±0.16<br>4.32% ** | 3.7±0.022<br>4.52% *** | 6.04±0.048<br>-1.15% |
| *aaBBdd* | 16 | 44.36±0.88<br>11.04% **** | 16.69±0.18<br>4.57% *** | 3.69±0.022<br>4.24% **** | 6.16±0.032<br>0.82% |
| *AAbbdd* | 30 | 44.79±0.4<br>12.12% **** | 16.83±0.12<br>5.45% **** | 3.68±0.016<br>3.95% **** | 6.21±0.029<br>1.64% ** |
| *aabbdd* | 37 | 46.33±0.38<br>15.97% **** | 17.24±0.1<br>8.02% **** | 3.74±0.013<br>5.65% **** | 6.28±0.026<br>2.78% **** |
| ANOVA (*P*-value) |  | < 2.2e-16 | 2.531e-15 | < 2.2e-16 | 3.175e-08 |

All data are shown as mean ± standard error. The phenotype data of each genotype was compared to that of genotype *AABBDD* using the two-tailed Student's t-test; \* significant at $0.01 < P < 0.05$ , \*\* significant at $0.001 < P < 0.01$ , \*\*\* significant at $0.0001 < P < 0.001$ , \*\*\*\* significant at $P < 0.0001$ . There were no phenotypic differences between re-generated plants with genotype *AABBDD* and wheat cultivar Bobwhite.

In Paragon, single mutants increased TGW by an average of 5.34%. However, differences between homoeologues were observed (unlike in Bobwhite), with only the *TaGW2-A1* and *TaGW2-B1* mutants resulting in significantly increased GS and TGW (*P* < 0.05; 8.6 and 9.5%, respectively), whereas the D-genome single mutant had no significant effects (−1.6%, Table 3 and Figure 2). The highest increase in GS and TGW in Paragon was associated with the KO mutation in the B genome (Table 3).-As in Bobwhite, no phenotypic differences between plants carrying wild type alleles in all three genomes (genotype *AABBDD*) and wheat cultivar Paragon were detected (except for grain length), suggesting that background mutations segregating in the progeny did not have substantial effects on the evaluated traits (Table 3).

**Figure 2.**
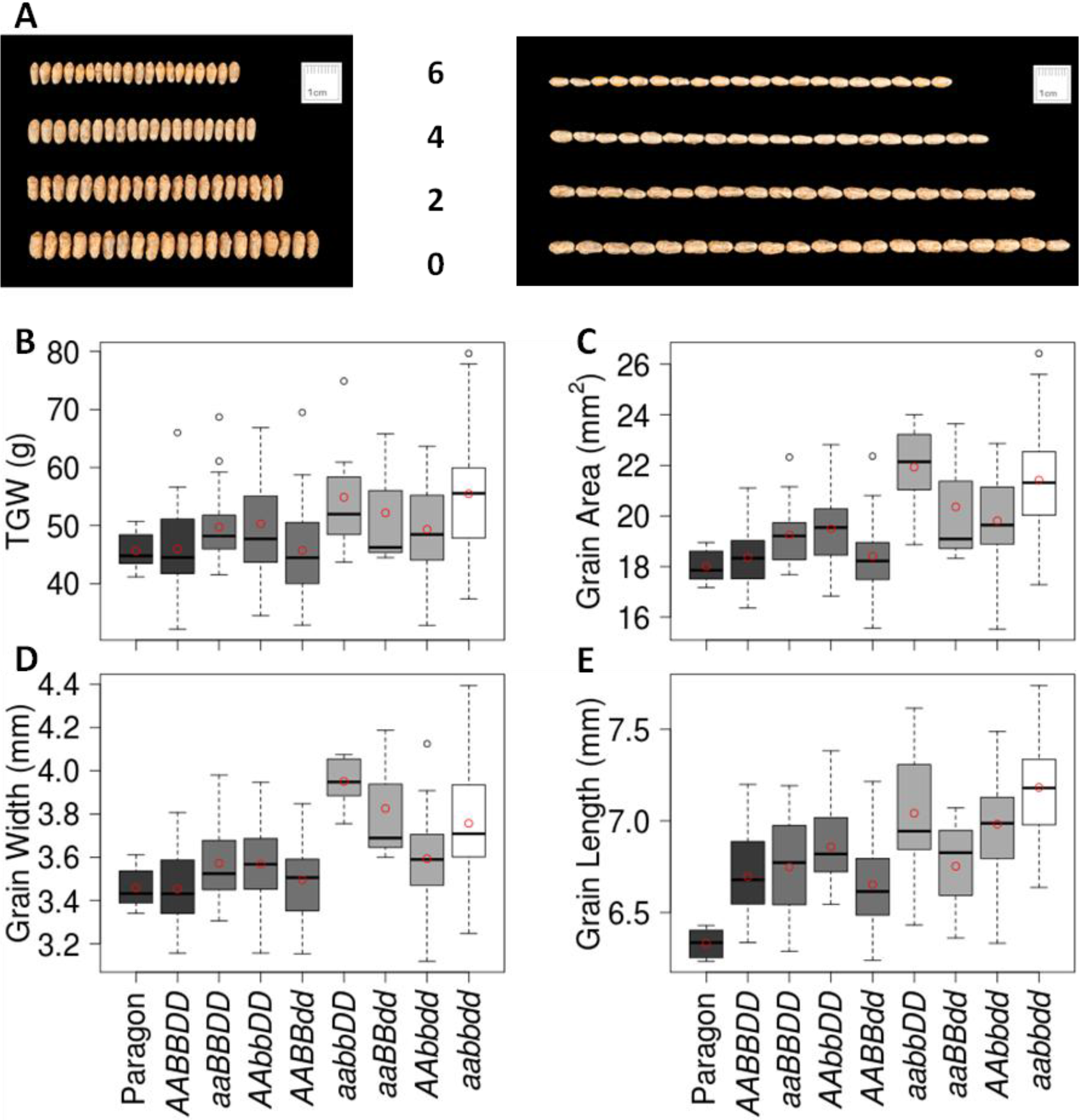
The effects of single-, double-, and triple-gene KO mutations in the *TaGW2* gene on the grain morphometric and TGW traits in Paragon. **A)** The image of twenty seeds from wild-type (6 functional alleles), single- (4), double- (2) and triple (0) mutant plants (scale bar 1 cm). **B-E)** Box and whisker plots show the distribution of TGW (B), grain area (C), grain width (D), and grain length (E) for wild-type and mutant wheat lines. The mean value for each genotype is shown as a red circle. The genotypes of the *TaGW2* homoeologues are similar to those in Figure 1.

**Table 3.** The TGW and grain morphometric parameters for the *TaGW2* gene mutants from the Paragon BC_1_F_3_ population.

| Genotype | *N* | TGW(g) | Area(mm<sup>2</sup>) | Width(mm) | Length(mm) |
| --- | --- | --- | --- | --- | --- |
| Paragon | 8 | 45.64±1.2 | 18.01±0.24 | 3.46±0.034 | 6.33±0.028<br>-5.52%**** |
| *AABBDD* | 32 | 45.98±1.21 | 18.36±0.2 | 3.45±0.03 | 6.7±0.039 |
| *aaBBDD* | 23 | 49.95±1.43<br>8.63%* | 19.33±0.25<br>5.23%** | 3.58±0.035<br>3.72%** | 6.78±0.054<br>1.19% |
| *AAbbDD* | 24 | 50.33±1.84<br>9.46%* | 19.5±0.31<br>6.20%** | 3.57±0.039<br>3.33%* | 6.86±0.041<br>2.45%** |
| *AABBdd* | 25 | 45.22±1.61<br>-1.65% | 18.27±0.28<br>-0.53% | 3.47±0.035<br>0.55% | 6.64±0.052<br>-0.86% |
| *aabbDD* | 7 | 54.89±3.95<br>19.38%** | 21.93±0.7<br>19.44%**** | 3.95±0.046<br>14.40%**** | 7.04±0.156<br>5.17%** |
| *aaBBdd* | 3 | 52.18±6.85<br>13.48% | 20.36±1.66<br>10.87%* | 3.83±0.183<br>10.80%** | 6.75±0.208<br>0.86% |
| *AAbbdd* | 22 | 49.34±1.73<br>7.31% | 19.81±0.38<br>7.90%*** | 3.59±0.047<br>4.06%* | 6.98±0.062<br>4.28%*** |
| *aabbdd* | 45 | 55.48±1.48<br>20.66%**** | 21.42±0.31<br>16.65%**** | 3.76±0.037<br>8.79%**** | 7.18±0.039<br>7.27%**** |
| ANOVA (*P* value) |  | 1.755e-05 | < 2.2e-16 | 8.395e-12 | < 2.2e-16 |

All data are shown as mean ± standard error. The phenotype data of each genotype and Paragon was compared to genotype AABBDD using the two-tailed Student's t-test, \* significant at 0.01 < *P* < 0.05, \*\* significant at 0.001 < *P* < 0.01, \*\*\* significant at 0.0001 < *P* < 0.001, \*\*\*\* significant at *P* < 0.0001. Except for grain length, there were no phenotypic differences between plants carrying wild type alleles in all three genomes (genotype AABBDD) and wheat cultivar Paragon.

### *TaGW2* displays a dosage-dependent effect on grain size and weight

To investigate whether the phenotypic effects of *TaGW2* were dosage-dependent, we evaluated double- and triple KO mutant lines of Bobwhite and Paragon. In Bobwhite, all double mutants (genotypes *aabbDD*, *aaBBdd*, and *AAbbdd*) had higher grain width and TGW compared to the respective single mutants (Table 1 and 2, Figure 1 and S4), with an average increase of 12.1% in TGW with respect to wild-type plants. These differences were significant (Student’s t-test *P* < 0.05) in at least one of the two experiments, except for the comparison of TGW between the A and D double mutant and the D genome single mutant lines (*P* = 0.23). Consistent with the B and D single mutants having higher contribution to GS/TGW than A single mutant, the grain length, grain area and TGW of the B and D double mutants were mostly the highest among all the double mutants except for grain width in one of the two experiments (Table 1 and 2, Figure 1 and S4).

In Paragon, the *aabbDD* and *aaBBdd* double mutants had a significant increase in grain width compared to the respective single mutants (Student’s t-test *P* < 0.05). Though not statistically significant, the TGW of *aabbDD* and *aaBBdd* double mutants was higher than the TGW of the respective single mutants (Table 3 and Figure 2). The grain area of the *aabbDD* double mutant was higher compared to the respective single mutants (Student’s t-test *P* < 0.05). Overall, double mutants increased TGW by an average of 10.53% with respect to wild-type plants.

The phenotypic effects of the *TaGW2* gene triple mutants were also investigated in both genetic backgrounds. In Bobwhite, the triple KO mutant had significantly higher GS and TGW compared to all three double mutants (Student’s t-test *P* < 0.05) in at least one experiment, except for the comparison of grain width between the triple and the *aabbDD* double mutant. On average, Bobwhite triple mutants increased TGW by 16.28%, grain width by 5.88% and grain length by 3.11% with respect to the wild type lines.

In Paragon, the distributions and mean values for GS and TGW in the triple mutants were close to those observed for double mutants in the A and B genomes (Student’s t-test *P* > 0.05), which was consistent with the lack of significant increase in GS and TGW in the single D-genome mutant (Figure 2 and Table 3). The Paragon triple mutant had significantly increased GS and TGW compared to both A and B single mutants (Student’s t-test *P* < 0.05), with an increase of 20.66% in TGW, 8.79% in grain width and 7.27% in grain length with respect to wild type lines.

The previous analyses were conducted with plants homozygous for mutations at each of the three genes. We further investigated the dosage-dependent response of grain morphometric traits and TGW by evaluating lines carrying the *TaGW2* gene mutations in homozygous or heterozygous states (Table S3). The analysis of GS and TGW traits in these lines showed that the distribution of GS and TGW negatively correlates with the number of the *TaGW2* functional alleles (Figure 3); the correlation coefficients for the TGW, grain area, grain width and grain length traits were −0.51, −0.38, −0.49, and −0.14, respectively. A similar relationship between phenotypic effects and the dosage of functional alleles was observed in the F_2_ population developed by crossing the Bobwhite triple mutant with cultivar Thatcher (Figure S5).

**Figure 3.**
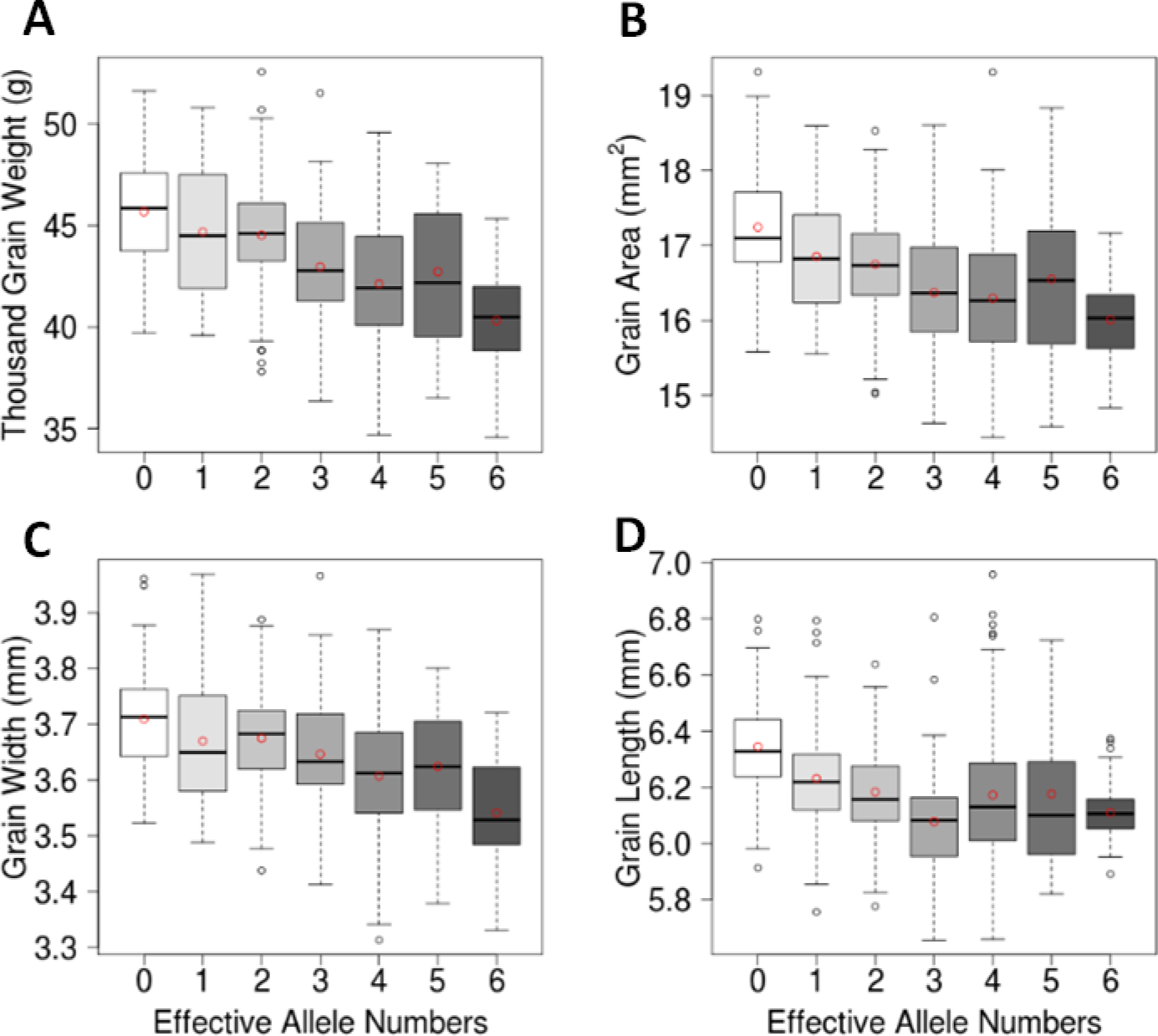
Relationship between the number of wild-type (non-mutant) *TaGW2* gene copies and grain morphometric and TGW traits in Bobwhite. Box and whisker plots show the trait distribution for TGW (A), grain area (B), grain width (C), and grain length (D) in Bobwhite gene edited mutants grouped based on the number of functional *TaGW2* copies. The mean value of each group is shown as a red circle within the box plot. The number of functional *TaGW2* copies is shown on the horizontal axis.

### Genome-specific bias and inter-cultivar variation in *TaGW2* homoeologue expression

A possible reason for the larger phenotypic effects of individual *TaGW2* homoeologues could be higher relative expression compared to the other gene copies. To test this hypothesis, we investigated the relative expression of *TaGW2-A1*, -*B1* and -*D1* in Bobwhite. In Bobwhite, *TaGW2-D1* had slightly higher expression level than *TaGW2-B1*, and the expression level of *TaGW2-A1* was significantly lower than that of the B and D genomes (Student’s t-test *P* <0.05) (Figure 4A). This expression pattern among the gene homoeologues was similar to that in Chinese Spring, where *TaGW2-A1* had the lowest and *TaGW2-D1* had the highest expression levels (Figure S6). The rank order of each homoeologue’s expression level in Bobwhite matched the rank order of the homoeologue’s KO effect on the GS and TGW traits.

**Figure 4.**
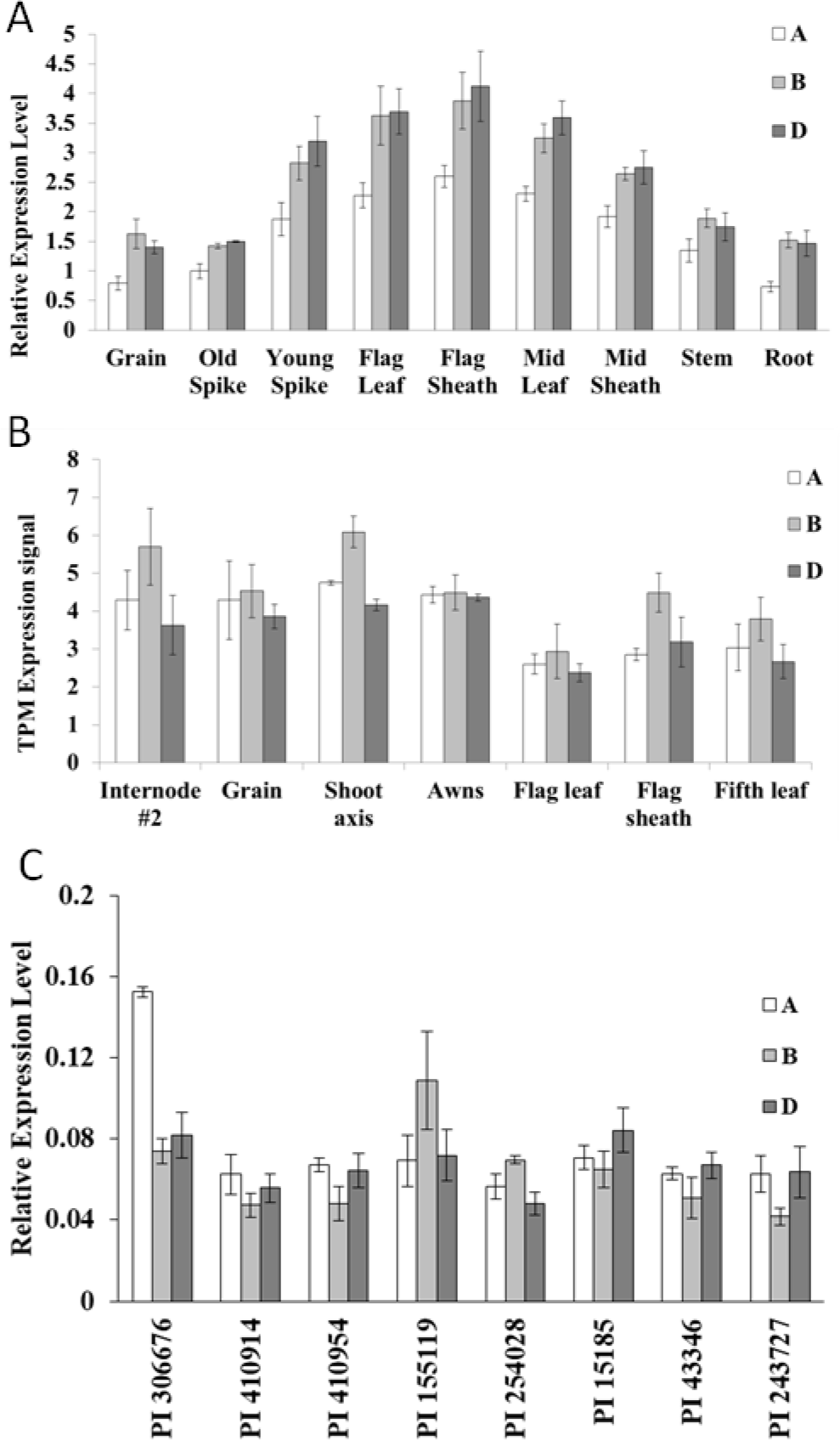
The expression patterns of the *TaGW2* gene homoeologues across different tissues and among different varieties. **A)** RT-PCR analysis of the relative expression levels of the three *TaGW2* homoeologues from different tissues of cultivar Bobwhite. **B)** Analysis of the relative expression levels of *TaGW2* homoeologue from different tissues of cultivar Azhurnaya using RNA-Seq data. Expression values are expressed in Transcripts Per Million (TPM) **C)** RT-PCR analysis of the relative expression levels of *TaGW2* homoeologues in a diverse panel of wheat lines using leaf tissues collected at two-leaf stage. Error bars denote standard error based on five biological and two technical replicates. The plant ID are from U.S. National Plant Germplasm System (https://npgsweb.ars-grin.gov/gringlobal/search.aspx?).

The expression patterns of *TaGW2-A1, -B1* and -*D1* described for Bobwhite and Chinese Spring were different from those reported for cultivars Shi4185 and Kenong99, both of which had the expression levels of three homoeologues ranked as follows: *TaGW2-B1* > *TaGW2-D1* > *TaGW2-A1* (Hong et al. 2014; Zhang et al. 2018). To understand whether the inter-cultivar variation in the contribution of the *TaGW2-A1, TaGW2-B1* and *TaGW2-D1* genes to total expression is a common phenomenon, we analyzed the *in-silico* expression browser of cultivar Azhurnaya and an additional eight wheat cultivars. Our results showed variation in the level of contribution of each homoeologue to the total expression of the *TaGW2* gene among these cultivars (Figure 4B and 4C). For example, in Azhurnaya the expression level of *TaGW2-B1* was significantly higher than that of *TaGW2-A1* and *TaGW2-D1*, and the expression of *TaGW2-D1* was only slightly lower than that of *TaGW2-A1* (Figure 4B). These results suggest the possibility that the grain morphometric and TGW traits in different cultivars are affected by the relative expression of the different genomic copies of *TaGW2*.

### Expression of the *TaGW2* homoeologues in the Bobwhite mutants

To investigate the effect of mutations on the expression of the *TaGW2* homoeologues, we assessed the expression levels of each *TaGW2* copy in single-, double-, and triple mutants of Bobwhite (Figure 5). Except for the A genome, the single mutants in the B and D genomes led to significant down-regulation of their own expression. However, the down-regulation of the *TaGW2-A1* gene expression was statistically significant only in the A/B genome double mutant. In all cases, mutations in a single gene copy did not result in significant compensatory changes in the expression of the other homoeologues. Only in the B/D double mutant there was a significant up-regulation of *TaGW2-A1* expression compared to the wild-type level (Student’s t-test *P* < 0.05). The remaining double mutants did not show changes in the expression of the remaining single functional copy compared to the wild-type plants.

**Figure 5.**
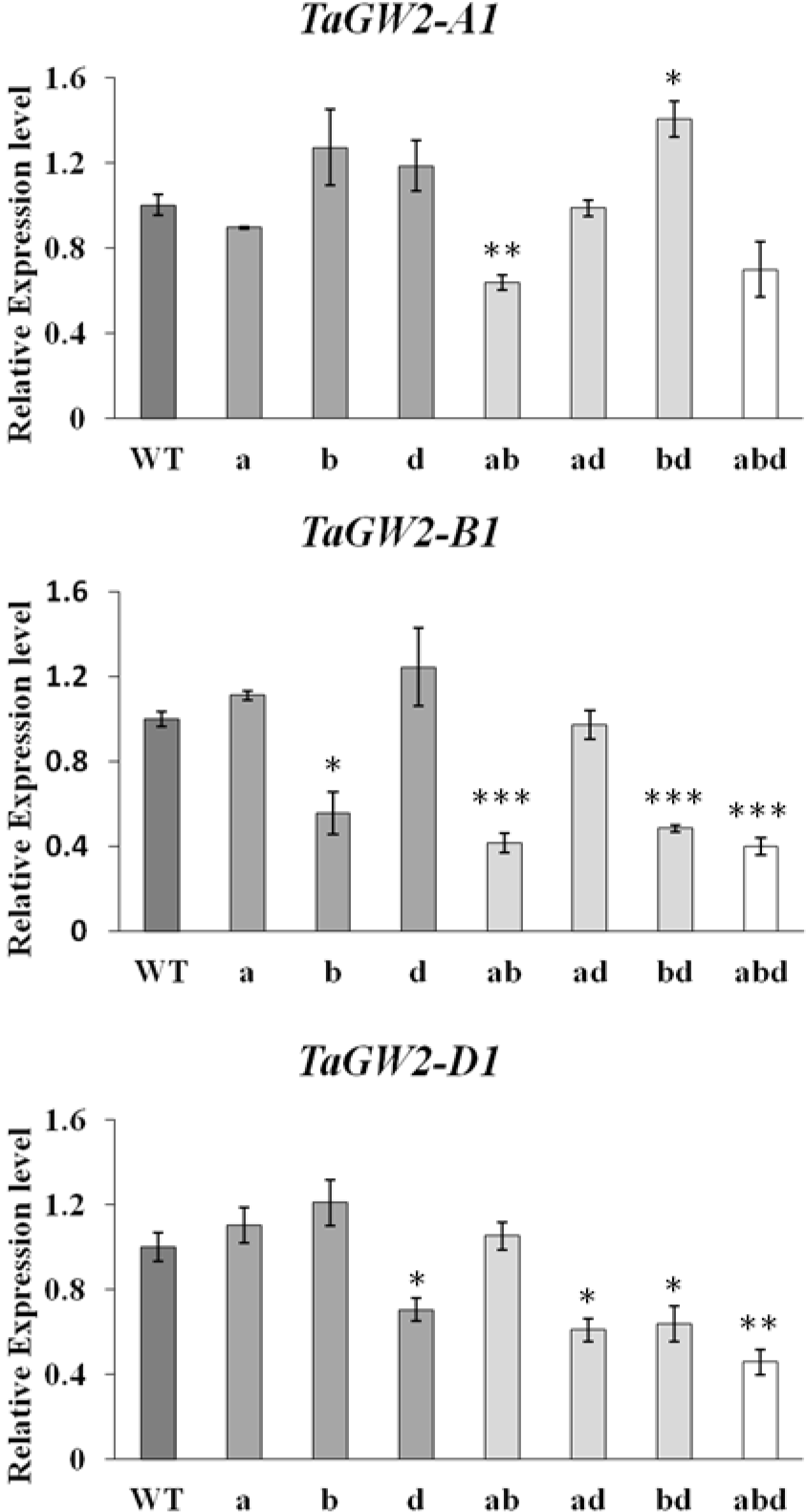
Transcript levels of the *TaGW2* homoeologues in wheat mutants. RT-PCR analysis of the *TaGW2* gene homoeologues using genome-specific primers (Fig. S3). The abbreviation WT, a, b, d, ab, ad, bd, and abd on the x-axis stand for genotypes *AABBDD*, *aaBBDD*, *AAbbDD*, *AABBdd*, *aabbDD*, *aaBBdd*, *AAbbdd*, and *aabbdd*, respectively. Expression values (mean±standard error based on three biological and two technical replicates) for each *TaGW2* homoeologue was first normalized using *actin* as internal control, and then shown as relative expression to the respective homoeologue in the wild-type plant. The relative expression levels of *TaGW2-A1, TaGW2-A1, and TaGW2-D1* in mutant plants were compared to that of the wild-type plant using the Student’s t-test; * significant at *P* < 0.05, ** significant at *P* < 0.01, *** significant at *P* < 0.001.

## Discussion

Using the wheat TILLING populations and the CRISPR-Cas9-based genome editing technology, we showed that the KO of each of the three homoeologues of the *TaGW2* gene, which are all functional in Bobwhite (Zhang et al. 2018), and the *A* and *B* genome copies of the *TaGW2* gene in Paragon increase GS and TGW. Our results are consistent with previous studies (Simmonds et al. 2016; Zhang et al. 2018) demonstrating that the functional copies of the *TaGW2* gene are negative regulators of GS and TGW. The only exception was the lack of significant effect of the *TaGW2-D1* gene KO in Paragon. However, the *aaBBdd* double mutants had a significant increase in grain width compared to the A and D genome single mutants, which indicated that the D genome single mutant might have effects on GS/TGW, albeit smaller compared to other genome copies.

Our study demonstrated that mutations in the homoeologous copies of the *TaGW2* gene have dosage-dependent effects on phenotype in both analyzed wheat cultivars. With the reduction in the number of functional gene copies, the grain morphometric traits and TGW tended to increase reaching maximum value in the lines with all *TaGW2* gene copies mutated. While in each cultivar one of the homoeologues showed more substantial effect than the others, most significant phenotypic changes were associated with changes in gene dosage rather than with particular combinations of mutated alleles. For example, in all possible double mutant combinations (AB, AD, BD), increase in GS and TGW was higher than in single-gene KO. These results were consistent with higher TGW in a mutant of cultivar Kenong99 that carried mutations in the B and D genome copies of the *TaGW2* gene compared to mutants carrying KO mutations in either B or D genomes (Zhang et al. 2018). These findings are consistent with previous studies suggesting the functional redundancy of duplicated genes in wheat (Borrill et al. 2015; Jordan et al. 2015; Uauy 2017), and indicate that by changing the dosages of functional homoeo-alleles it should be possible to expand the range of phenotypic variation in wheat available for trait improvement.

Previously, it was suggested that *TaGW2-B1* might be functionally more active than *TaGW2-D1* (Zhang et al. 2018). Consistent with this finding, in Paragon, the KO of *TaGW2-B1* had larger effects on some grain morphometric traits compared to the mutants of *TaGW2-A1* or *TaGW2-D1*. However, the finding that the mutation with the highest effect in Bobwhite is located in the D genome suggests the presence of inter-cultivar variation for the contribution of *TaGW2-A1, TaGW2-B1* and *TaGW2-D1* to regulate yield component traits. One possible factor underlying this variation can be inter-cultivar differences in the contribution of different homoeologues to the total level of *TaGW2* gene expression. Our data is consistent with this hypothesis by showing that the KO of the homoeologue expressed at the highest level had the highest effect on phenotype. The lack of direct effect of homoeologue KOs on the expression of other non-mutated gene copies, also suggests that the total level of the functional *TaGW2* gene transcripts is defined by the dose of the non-mutated homoeologues. Likewise, previous studies demonstrated that the level of *TaGW2* gene expression is negatively related to the GS and TGW (Hong et al. 2014; Qin et al. 2014; Qin et al. 2017; Su et al. 2011). The analysis of a diverse set of wheat cultivars corroborated that inter-cultivar variation in the relative expression of the *TaGW2* homoeologues is common in wheat and suggest that variation in the contribution of different genomes to gene expression might play important roles in phenotypic variation in wheat. Understanding the basis of this homoeologue-specific expression bias in natural populations will be critical for connecting phenotype with genotype in future studies. For the *TaGW2* gene, extensive difference among the homoeologous copies were discovered in the promoter regions (Qin et al. 2017), suggesting that diversity in the regulatory sequences can be one of the factors driving inter-cultivar expressional diversity in this gene.

In the previous study, the triple-mutants of both Kenong99 and Bobwhite produced wrinkled grains (Zhang et al. 2018). Likewise, in our study, wrinkled grains were found in the double- (A and B genomes) and triple-mutants of Paragon. It was speculated that wrinkled grains in the triple mutants might result from significant reductions of the *TaGW2* function (Zhang et al. 2018). However, no wrinkled grains were found in the triple-gene KO mutants of Bobwhite in our study, which suggest this effect might result from insufficient grain filling under glasshouse conditions. The total KO of *TaGW2* gene function could strongly increase the maternal pericarp cell growth, and thus increase the space for grain filling (Simmonds et al. 2016; Zhang et al. 2018). If the source is unable to fill grain due to low productivity, or reduced efficiency of transport from source to sink, the enlarged seeds on the *TaGW2* gene mutants could be wrinkled after maturation. It is possible that the deficient grain filling could be offset by optimal growth conditions promoting active grain filling under field conditions or by combining the trait with high biomass lines.

## Conclusion

By using TILLING and the CRISPR-Cas9-based genome editing strategy, we demonstrated that all three homoeologues copies of the *TaGW2* gene act as negative regulators of GS and TGW. By combining homoeologous KO mutations in *TaGW2* in all possible combinations, we showed that homoeologues act additively imposing dosage-dependent effects on both total gene expression and phenotype. The newly generated alleles of *TaGW2* will allow combining mutant alleles in different configurations which should allow fine tuning of the phenotypic effects. The marker-assisted selection efforts will be facilitated by the availability of previously reported genome-specific Cleaved Amplified Polymorphic Sequences markers for the CRISPR-Cas9-induced mutations (Wang et al. 2018), as well as KASP markers that can be developed for the EMS-induced mutations in this study.

Here, we also report cultivar-specific phenotypic effects for different homoeologous copies of the *TaGW2* gene. This inter-cultivar variation appears to be associated with the level of homoeologue contribution to the total level of *TaGW2* gene expression. The results from the expression study across varieties suggests that understanding the relative contribution of each homoeologue in the target varieties will be an important considering and should help determine the best mutant combinations to deploy to improve grain size, and potentially yields in the field. Taken together, our results indicate that the polyploid origin of the wheat genome provides ample opportunities for diversifying the genetic architecture of agronomic traits. The targeted mutagenesis of allopolyploid genome on genes with the potential to affect major traits can be a powerful tool for expanding existing allelic variation available for wheat improvement.

## Author contribution statement

WW – designed and conducted gene editing experiments, generated and analyzed gene expression data, developed F2 population, drafted the manuscript; JS – developed wheat lines with EMS KOs of the *TaGW2* gene, collected and analyzed data, contributed to preparing the manuscript; QP – analyzed gene editing events using next generation sequencing (NGS) and conducted *TaGW2* gene expression analysis; FH – wrote scripts for the NGS analysis of editing events and analyzed data; AB – identified the EMS-induced B genome mutations in the *TaGW2* gene; AA – designed experiments for NGS analysis of editing events and performed NGS; HT – performed biolistic transformation of wheat embryos with the gene editing constructs; CU – conceived idea, analyzed data and contributed to manuscript writing; EA – conceived idea, designed gene editing experiments, coordinated project, analyzed data and wrote the manuscript.

## Acknowledgements

This project was supported by the National Research Initiative Competitive Grants 2017-67007-25932 from the USDA National Institute of Food and Agriculture, the International Wheat Yield Partnership grant IWYP76, the Bill and Melinda Gates Foundation grant BMGF:01511000146, and the UK Biotechnology and Biological Sciences Research Council (BBSRC) Designing Future Wheat (BB/P016855/1) and GEN (BB/P013511/1) programs.

## References

Avni R, Nave M, Barad O, Baruch K, Twardziok SO, Gundlach H, Hale I, Mascher M, Spannagl M, Wiebe K, Jordan KW, Golan G, Deek J, Ben-Zvi B, Ben-Zvi G, Himmelbach A, MacLachlan RP, Sharpe AG, Fritz A, Ben-David R, Budak H, Fahima T, Korol A, Faris JD, Hernandez A, Mikel MA, Levy AA, Steffenson B, Maccaferri M, Tuberosa R, Cattivelli L, Faccioli P, Ceriotti A, Kashkush K, Pourkheirandish M, Komatsuda T, Eilam T, Sela H, Sharon A, Ohad N, Chamovitz DA, Mayer KFX, Stein N, Ronen G, Peleg Z, Pozniak CJ, Akhunov ED, Distelfeld A (2017) Wild emmer genome architecture and diversity elucidate wheat evolution and domestication. Science 357:93–96

Borrill P, Adamski N, Uauy C (2015) Genomics as the key to unlocking the polyploid potential of wheat. New Phytol 208:1008–1022

Borrill P, Ramirez-Gonzalez R, Uauy C (2016) expVIP: a Customizable RNA-seq Data Analysis and Visualization Platform. Plant Physiol 170:2172–2186

Cavanagh CR, Chao S, Wang S, Huang BE, Stephen S, Kiani S, Forrest K, Saintenac C, Brown-Guedira GL, Akhunova A, See D, Bai G, Pumphrey M, Tomar L, Wong D, Kong S, Reynolds M, da Silva ML, Bockelman H, Talbert L, Anderson JA, Dreisigacker S, Baenziger S, Carter A, Korzun V, Morrell PL, Dubcovsky J, Morell MK, Sorrells ME, Hayden MJ, Akhunov E (2013) Genome-wide comparative diversity uncovers multiple targets of selection for improvement in hexaploid wheat landraces and cultivars. Proc Natl Acad Sci U S A 110:8057–8062

Choulet F, Alberti A, Theil S, Glover N, Barbe V, Daron J, Pingault L, Sourdille P, Couloux A, Paux E, Leroy P, Mangenot S, Guilhot N, Le Gouis J, Balfourier F, Alaux M, Jamilloux V, Poulain J, Durand C, Bellec A, Gaspin C, Safar J, Dolezel J, Rogers J, Vandepoele K, Aury JM, Mayer K, Berges H, Quesneville H, Wincker P, Feuillet C (2014) Structural and functional partitioning of bread wheat chromosome 3B. Science 345:1249721

Cong L, Ran FA, Cox D, Lin S, Barretto R, Habib N, Hsu PD, Wu X, Jiang W, Marraffini LA, Zhang F (2013) Multiplex genome engineering using CRISPR/Cas systems. Science 339:819–823

Hong Y, Chen L, Du LP, Su Z, Wang J, Ye X, Qi L, Zhang Z (2014) Transcript suppression of TaGW2 increased grain width and weight in bread wheat. Funct Integr Genomics 14:341–349

Jaiswal V, Gahlaut V, Mathur S, Agarwal P, Khandelwal MK, Khurana JP, Tyagi AK, Balyan HS, Gupta PK (2015) Identification of Novel SNP in Promoter Sequence of TaGW2-6A Associated with Grain Weight and Other Agronomic Traits in Wheat (Triticum aestivum L.). PLoS One 10:e0129400

Jinek M, Chylinski K, Fonfara I, Hauer M, Doudna JA, Charpentier E (2012) A programmable dual-RNA-guided DNA endonuclease in adaptive bacterial immunity. Science 337:816–821

Jordan KW, Wang S, Lun Y, Gardiner LJ, MacLachlan R, Hucl P, Wiebe K, Wong D, Forrest KL, Consortium I, Sharpe AG, Sidebottom CH, Hall N, Toomajian C, Close T, Dubcovsky J, Akhunova A, Talbert L, Bansal UK, Bariana HS, Hayden MJ, Pozniak C, Jeddeloh JA, Hall A, Akhunov E (2015) A haplotype map of allohexaploid wheat reveals distinct patterns of selection on homoeologous genomes. Genome Biol 16:48

Krasileva KV, Vasquez-Gross HA, Howell T, Bailey P, Paraiso F, Clissold L, Simmonds J, Ramirez-Gonzalez RH, Wang X, Borrill P, Fosker C, Ayling S, Phillips AL, Uauy C, Dubcovsky J (2017) Uncovering hidden variation in polyploid wheat. Proc Natl Acad Sci U S A 114:E913–E921

Li W, Yang B (2017) Translational genomics of grain size regulation in wheat. Theor Appl Genet 130:1765–1771

Liang Z, Chen K, Li T, Zhang Y, Wang Y, Zhao Q, Liu J, Zhang H, Liu C, Ran Y, Gao C (2017) Efficient DNA-free genome editing of bread wheat using CRISPR/Cas9 ribonucleoprotein complexes. Nat Commun 8:14261

Mali P, Yang L, Esvelt KM, Aach J, Guell M, DiCarlo JE, Norville JE, Church GM (2013) RNA-guided human genome engineering via Cas9. Science 339:823–826

Minkenberg B, Wheatley M, Yang Y (2017) CRISPR/Cas9-Enabled Multiplex Genome Editing and Its Application. Prog Mol Biol Transl Sci 149:111–132

Mohan C (2016) Genome Editing in Sugarcane: Challenges Ahead. Front Plant Sci 7:1542

Pearce S, Vazquez-Gross H, Herin SY, Hane D, Wang Y, Gu YQ, Dubcovsky J (2015) WheatExp: an RNA-seq expression database for polyploid wheat. BMC Plant Biol 15:299

Qin L, Hao C, Hou J, Wang Y, Li T, Wang L, Ma Z, Zhang X (2014) Homologous haplotypes, expression, genetic effects and geographic distribution of the wheat yield gene TaGW2. BMC Plant Biol 14:107

Qin L, Zhao J, Li T, Hou J, Zhang X, Hao C (2017) TaGW2, a Good Reflection of Wheat Polyploidization and Evolution. Front Plant Sci 8:318

Ramirez-Gonzalez RH, Uauy C, Caccamo M (2015) PolyMarker: A fast polyploid primer design pipeline. Bioinformatics 31:2038–2039

Simmonds J, Scott P, Brinton J, Mestre TC, Bush M, Del Blanco A, Dubcovsky J, Uauy C (2016) A splice acceptor site mutation in TaGW2-A1 increases thousand grain weight in tetraploid and hexaploid wheat through wider and longer grains. Theor Appl Genet 129:1099–1112

Song XJ, Huang W, Shi M, Zhu MZ, Lin HX (2007) A QTL for rice grain width and weight encodes a previously unknown RING-type E3 ubiquitin ligase. Nat Genet 39:623–630

Su Z, Hao C, Wang L, Dong Y, Zhang X (2011) Identification and development of a functional marker of TaGW2 associated with grain weight in bread wheat (Triticum aestivum L.). Theor Appl Genet 122:211–223

Uauy C (2017) Wheat genomics comes of age. Curr Opin Plant Biol 36:142–148

Uauy C, Paraiso F, Colasuonno P, Tran RK, Tsai H, Berardi S, Comai L, Dubcovsky J (2009) A modified TILLING approach to detect induced mutations in tetraploid and hexaploid wheat. BMC Plant Biol 9:115

Wang SC, Wong DB, Forrest K, Allen A, Chao SM, Huang BE, Maccaferri M, Salvi S, Milner SG, Cattivelli L, Mastrangelo AM, Whan A, Stephen S, Barker G, Wieseke R, Plieske J, Lillemo M, Mather D, Appels R, Dolferus R, Brown-Guedira G, Korol A, Akhunova AR, Feuillet C, Salse J, Morgante M, Pozniak C, Luo MC, Dvorak J, Morell M, Dubcovsky J, Ganal M, Tuberosa R, Lawley C, Mikoulitch I, Cavanagh C, Edwards KJ, Hayden M, Akhunov E, Sequencing IWG (2014a) Characterization of polyploid wheat genomic diversity using a high-density 90 000 single nucleotide polymorphism array. Plant Biotechnol J 12:787–796

Wang W, Akhunova A, Chao S, Akhunov E (2016) Optimizing multiplex CRISPR/Cas9-based genome editing for wheat. BioRxiv

Wang W, Pan Q, He F, Akhunova A, Chao S, Trick H, Akhunov E (2018) Transgenerational CRISPR-Cas9 Activity Facilitates Multiplex Gene Editing in Allopolyploid Wheat. The CRISPR Journal 1:65–74

Wang Y, Cheng X, Shan Q, Zhang Y, Liu J, Gao C, Qiu JL (2014b) Simultaneous editing of three homoeoalleles in hexaploid bread wheat confers heritable resistance to powdery mildew. Nat Biotechnol 32:947–951

Yang Z, Bai Z, Li X, Wang P, Wu Q, Yang L, Li L, Li X (2012) SNP identification and allelic-specific PCR markers development for TaGW2, a gene linked to wheat kernel weight. Theor Appl Genet 125:1057–1068

Zadoks J, Chang T, Konzak C (1974) A decimal code for the growth stages of cereals. Weed Research 14:415–421

Zhang Y, Li D, Zhang D, Zhao X, Cao X, Dong L, Liu J, Chen K, Zhang H, Gao C, Wang D (2018) Analysis of the functions of TaGW2 homoeologs in wheat grain weight and protein content traits. Plant J

Zhang Y, Liang Z, Zong Y, Wang Y, Liu J, Chen K, Qiu JL, Gao C (2016) Efficient and transgene-free genome editing in wheat through transient expression of CRISPR/Cas9 DNA or RNA. Nat Commun 7:12617

